# Morning sickness-like changes during pregnancy in non-human primates and rodents

**DOI:** 10.1101/2025.08.14.668061

**Authors:** Saori Yano-Nashimoto, Kazutaka Shinozuka, Takuma Kurachi, Katsura Kagawa, Kentaro Q. Sakamoto, Yuko Shigeno, Kimie Niimi, Kumi O. Kuroda, Soichiro Yamaguchi

**Author notes:** **Corresponding author:** Saori Yano-Nashimoto, Laboratory of Physiology, Department of Basic Veterinary Sciences, Faculty of Veterinary Medicine, Hokkaido University, Kita 18, Nishi 9, Kita-ku, Sapporo, Hokkaido, 060-0818, Japan.

## Abstract

Nausea and vomiting of pregnancy (NVP), commonly referred to as “morning sickness,” causes a range of symptoms including nausea, vomiting, weight loss, anorexia, and malaise in pregnant women during early pregnancy. Although NVP highly affects the quality of life of pregnant women, the absence of model animals has prevented us from revealing the underlying mechanisms. Here, to identify whether non-human animals may serve as models for studying NVP, we analyzed metabolic changes during pregnancy in common marmosets and mice. Marmosets exhibited a transient weight decrease during the period of placental development in approximately 22% of pregnancies. Some marmosets repeatedly showed transient weight loss across multiple pregnancies, suggesting individual variations in the likelihood of NVP-like symptoms. Although mice did not show apparent alteration in body weight, they exhibited a slowdown in food intake and alterations in locomotor activity during the corresponding phase. These shared alterations suggest that marmosets and mice may exhibit phenomena analogous to human NVP and could serve as useful models for investigating its biological basis.

**Highlights:**

- Marmosets exhibited a transient weight loss during placental development in 22% of pregnancies.
- There was individual variation in how often the transient weight loss occurred in marmosets.
- Food intake and activity in C57BL/6 mice are altered during the corresponding gestational phase.
- Multiple morning sickness-like phenomena were observed in non-human mammals.

## Introduction

Nausea and vomiting of pregnancy (NVP), commonly referred to as “morning sickness”, causes a wide range of symptoms including nausea, vomiting, anorexia, weight decrease, malaise, and alterations in taste and olfaction in pregnant women. These symptoms are typically most severe during the 1^st^ trimester, when placental development is most active. NVP affects approximately 70-80% of pregnant women and impairs their quality of life^1–3^. In addition, in severe cases, several studies consistently find a general association between NVP and preterm birth, low birthweight, and fetal growth restriction, as well as some evidence of longer-term detrimental impacts on offspring such as neurodevelopmental delay^4^. However, the underlying mechanisms remain poorly understood, and thus no definitive treatment currently exists^2,5^.

One major obstacle to elucidating the mechanisms of NVP is the lack of appropriate non-human animal models. Several mammalian species have been reported to show changes reminiscent of human NVP^6,7^. For example, dogs may show reduced food intake and occasional vomiting during early pregnancy^6^. Pregnant rhesus monkeys have been observed to reject food more frequently^7^. However, because these observations have not been followed by systematic scientific investigation, it is unclear whether these alterations can lead to more severe symptoms such as weight decrease, and whether the timing corresponds to the gestational stage of human NVP. Thus, such phenomena are not widely recognized as equivalent to human NVP, and these species are not considered animal models for studying NVP.

Here, to identify a potential non-human model for investigating the mechanisms of NVP, we carefully analyzed metabolic changes during pregnancy in common marmosets (*Callithrix jacchus*) and mice (*Mus musculus*), widely used experimental animals. As direct evaluation of nausea and vomiting is challenging in mice and in marmosets housed in family groups, we instead monitored physiological and behavioral indices such as body weight, food intake, and locomotor activity, which are often associated with nausea and malaise in the context of sickness behavior^8–10^. We found that some individuals in marmosets exhibited a transient decrease in body weight during the period of placental development. Furthermore, in mice, we observed a transient slowdown in food intake and alterations in locomotor activity during the corresponding gestational phase. These shared alterations suggest that both species may exhibit phenomena analogous to human NVP.

## Results

### Transient weight loss during mid-pregnancy was revealed by cluster analysis in pregnant marmosets

We conducted a cluster analysis on the body weight changes in pregnant marmosets from two different laboratories: Laboratory K (6 mothers, 27 pregnancies) and Laboratory N (14 mothers, 88 pregnancies) (Fig. 1A-B). The data from these laboratories were analyzed separately because the frequency of weight measurements differed. Hierarchical clustering was performed based on pairwise correlation dissimilarities. The data were grouped into three clusters by cutting the dendrogram (Fig. 1C-H). In Laboratory K, 13 pregnancies were classified as cluster A, which showed a consistent increase in body weight (Fig. 1C, E, G, I). Twelve and two pregnancies were classified as cluster B and C, respectively, both of which exhibited a transient decrease during mid-pregnancy, approximately 95-65 days before delivery. Similarly, in Laboratory N, the largest cluster (Cluster 1) showed a consistent increase, while the second cluster (Cluster 2) showed a transient decrease during mid-pregnancy (Fig. 1D, F, H, I). Notably, the timing of weight decrease was strikingly similar across Cluster B, C, and 2 (Fig. 1G, H), and this period corresponds to the phase of placental development in marmosets^11,12^.

**Figure 1:**
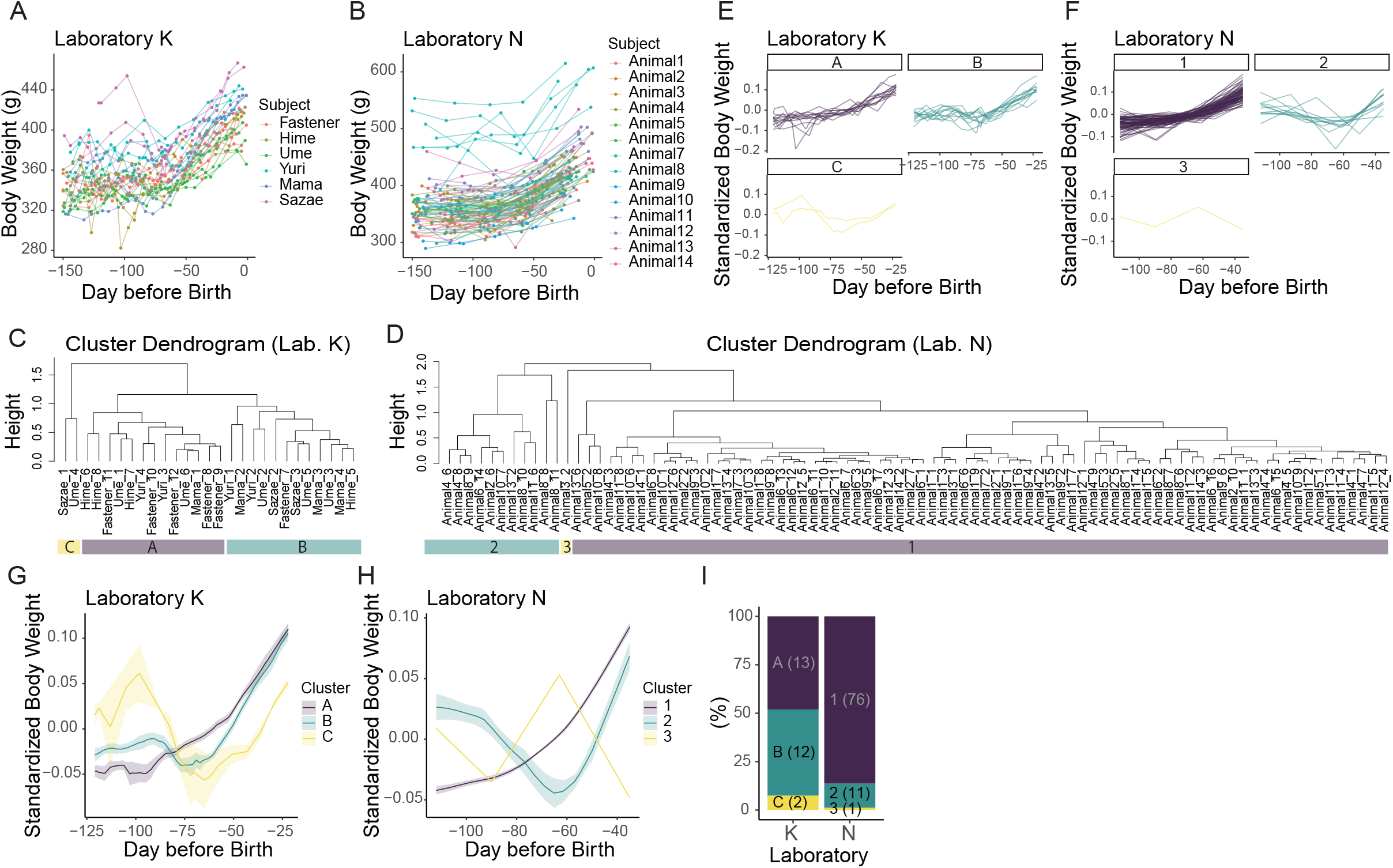
Clustering of body weight changes in pregnant marmosets. A-B Body weight changes in pregnant marmosets from two laboratories (Laboratory K and N). The x-axis shows days before birth (Day 0). C-D Cluster dendrograms based on pairwise correlations of body weight trajectories during pregnancy. Each leaf represents a pregnancy. The height of each node corresponds to the dissimilarity (1 – correlation coefficient) at which clusters are joined. Cluster labels (A–C or 1-3) are indicated below the leaves. E-H Body weight changes during pregnancy, shown separately for each cluster. For each pregnancy, body weight was standardized by its mean value. In panels E and F, each line represents a pregnancy. In panels G and H, the mean ± standard error is shown in each cluster. I The percentage of each cluster in each laboratory. The numbers within parentheses are the numbers of pregnancies included in the cluster.

Given the similarity in cluster characteristics across the two laboratories, we combined the data and re-classified them into two superclusters: Decrease-No and Decrease-Yes. Decrease-No includes Cluster A and 1, while Decrease-Yes includes Cluster B, C, and 2 (Fig. 2A, B). Cluster 3, which included only one pregnancy, was classified as “Other”. From 95 to 65 days before birth, body weight in the Decrease-No group increased by 3.4 ± 0.3% relative to the average pregnancy weight. In contrast, the Decrease-Yes group showed a decrease of 6.3 ± 1.5% during the same period, followed by a subsequent increase (Fig. 2A). There was individual variation in how often transient weight loss occurred during mid-pregnancy (Fig. 2B-F). Although the proportion of Decrease-Yes pregnancies differed between Laboratory K and N (Fig. 1I, 2B), some of the mothers showed this pattern repeatedly (Fig. 2D, F), whereas others rarely did (Fig. 2C, E).

**Figure 2:**
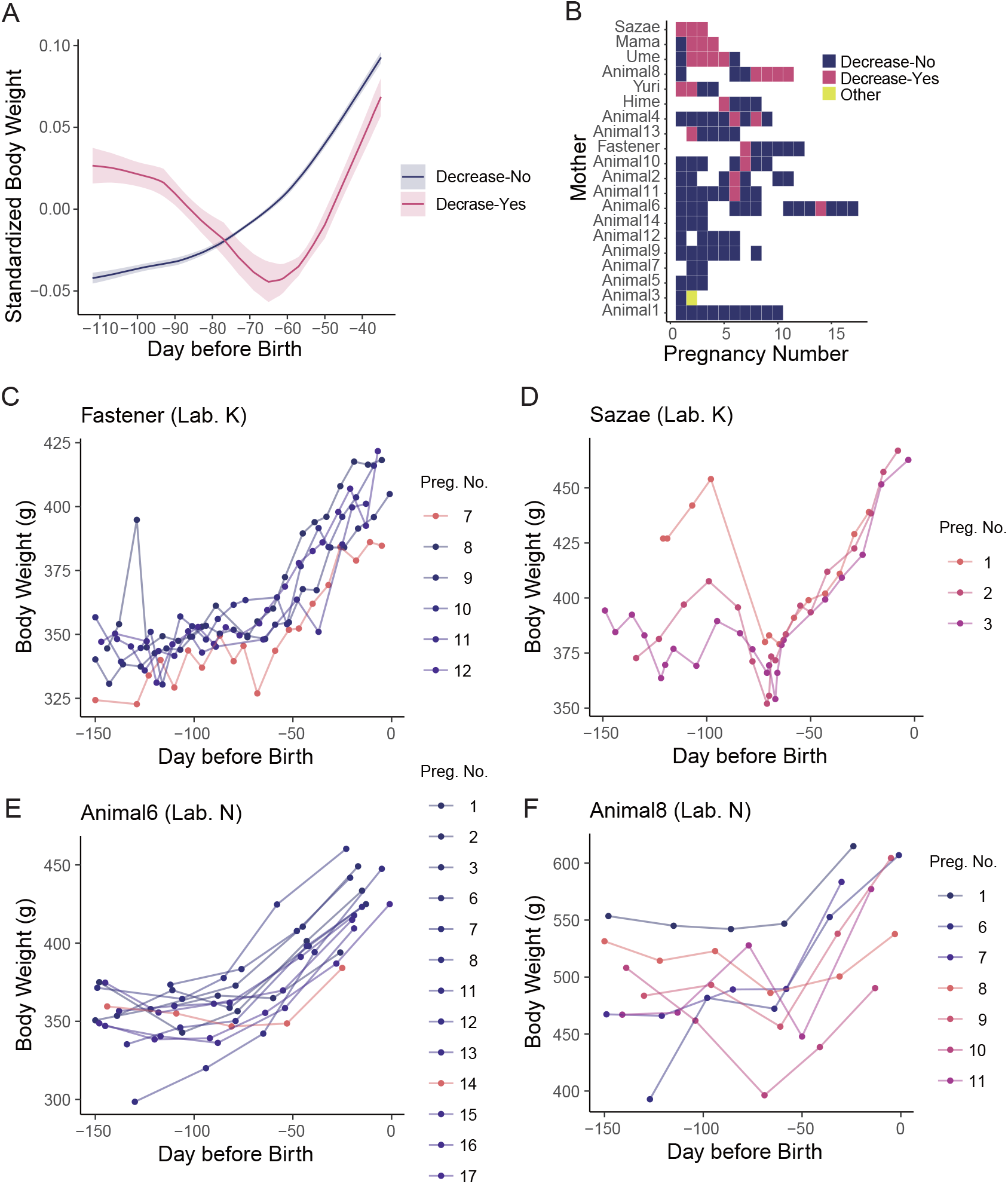
A subset of marmoset mothers repeatedly exhibited transient decreases in body weight during mid-pregnancy. A Body weight changes during pregnancy, shown separately for two superclusters: “Decrease-No”, and “Decrease-Yes”. “Decrease-No” includes Cluster A and Cluster 1; “Decrease-Yes” includes Cluster B, C, and 2. Cluster 3 is classified as “Other” and not shown in this panel. Body weight was standardized by its mean value. The mean ± standard error is shown in each cluster. B Tile plot showing cluster classification for each mother and pregnancy. Colors indicate cluster membership. C-F Body weight changes during pregnancy in representative mothers. The colors indicate cluster classification: red for “Decrease-Yes” and blue for “Decrease-No.” Panels C and E show mothers who were less likely to experience weight loss during pregnancy. Panels D and F show mothers who frequently lost weight during pregnancy. Mothers in panels C and D are from Laboratory K, and those in panels E and F are from Laboratory N.

### Mice showed transient alterations in food intake and locomotion during mid-pregnancy

To examine whether similar alterations occur in mice, we measured the body weight, food intake, and locomotor activity during pregnancy. Morning sickness in humans and transient weight decrease in marmosets are observed during the phase of placental development. Therefore, we compared these indices during the second quarter of pregnancy (Q2), when placentation progresses^13^, with those in other gestational phases. Body weight increased throughout pregnancy and showed no apparent alteration during Q2 (Fig. 3A). Meanwhile, although the food intake increased throughout pregnancy, the rate of increase slowed during Q2 (Fig. 3B). Locomotor activity increased according to the progress of gestation during the first quarter of pregnancy, plateaued during Q2, and then declined during the third quarter, when weight gain became more pronounced (Fig. 3A, C). The slowdown of the increase in locomotor activity during Q2 may suggest mild anorexia^9,10^. These results suggest that mice also exhibit metabolic alterations during the phase of placental development, as seen in marmosets and humans, although the changes appear to be more subtle.

**Figure 3:**
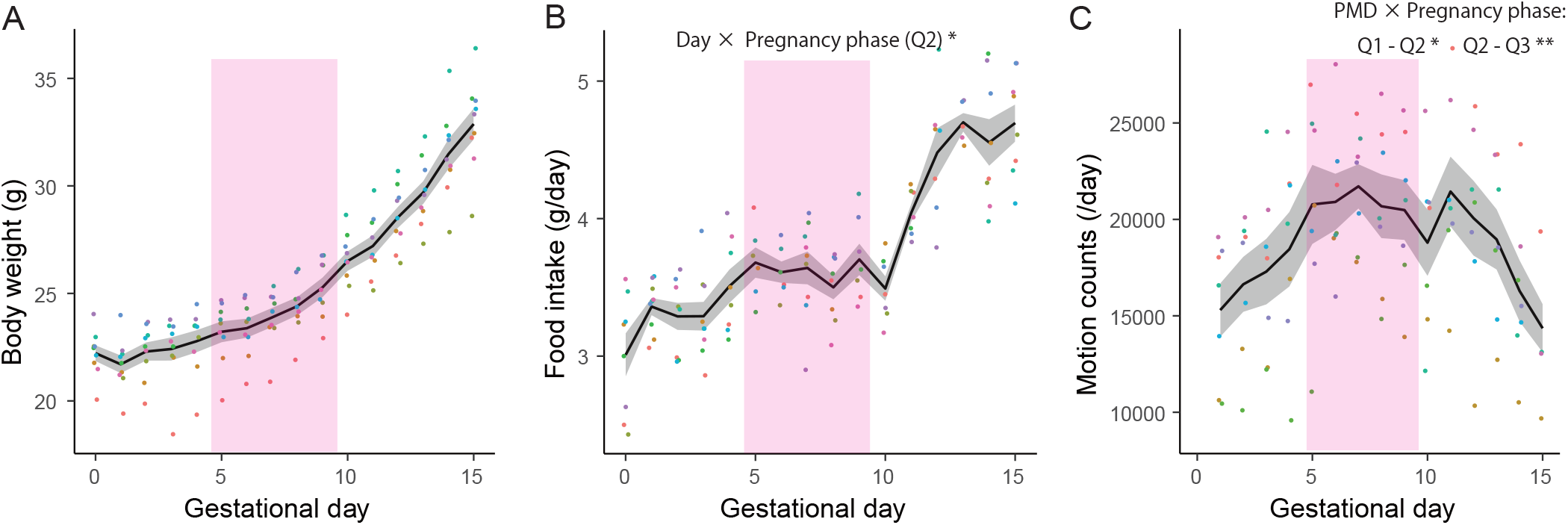
Mice exhibited transient alterations in food intake and locomotion during mid-pregnancy. A Body weight changes during pregnancy. Body weight increased throughout pregnancy. B Changes in food intake during pregnancy. Although food intake increased throughout pregnancy, the rate of increase slowed during the second quarter of pregnancy (Q2). *LMM, *p* < 0.05. C Changes in locomotor activity during pregnancy. Locomotor activity increased according to the progress of gestation during the first quarter of pregnancy, plateaued during Q2, and then declined during the third quarter. LMM, \*\**p* < 0.01, \**p* < 0.05. The mean ± standard error is shown as a black line with a gray shaded area. The color of each dot represents an individual mouse. Pink shading indicates Q2 period (gestational day 5 – 9).

## Discussion

We found that non-human mammals also showed NVP-like alteration during the period when placentation progresses^11–13^. In marmosets, a transient decrease in body weight was observed in a subset of pregnancies. In mice, although no body weight decrease was observed, the increase in food intake and locomotor activity slowed during the same gestational period, suggesting the possibility of mild anorexia and malaise. The possibility of NVP-like phenomena in non-human mammals has been suggested anecdotally, often based on isolated reports from zookeepers or animal caretakers. However, scientific investigations remain limited, and the few existing studies have not provided strong evidence that these phenomena are equivalent to human NVP^6,7^. In this study, we identified similar alterations occurring at a stage of pregnancy in marmosets and mice that corresponded to the timing of NVP in humans^11–14^. This provides novel and compelling evidence that the physiological basis of NVP may be evolutionarily conserved among mammals.

Among the diverse symptoms associated with NVP, we observed weight decrease, anorexia, and malaise-like alterations in mice or marmosets. However, the most famous symptom in NVP, vomiting^15,16^, cannot be assessed in the present study because mice do not vomit^8^ and because our marmoset subjects were kept in the family. Since marmosets sometimes vomit, it may be of interest to investigate its occurrence in pregnant marmosets in future research. Alterations in taste and odor in pregnant animals were reported in some research^17–19^, but the timing does not fully correspond to the gestational stage of human NVP. Thus, it should be carefully considered whether these alterations were related to NVP. In addition, given the diverse symptoms of NVP, it is possible that different symptoms are driven by distinct mechanisms (Anan et al, under review), underscoring the need for symptom-specific investigations.

In marmosets, transient weight decreases were observed only in approximately 22% of pregnancies, rather than universally. This proportion is comparable to the reported prevalence of moderate to severe NVP (with vomiting) in humans^3^. Similar to the variation in NVP susceptibility among women^20^, some marmoset females repeatedly exhibited transient weight loss across multiple pregnancies, suggesting the possibility of individual variations in the likelihood and severity of NVP-like symptoms. In mice, such variation was not observed, possibly due to their uniform genetic background. While previous studies have examined changes in food intake and activity levels during pregnancy in mice^21,22^, NVP-like symptoms have received little attention. This may be because the changes are relatively mild and therefore difficult to detect. In addition, differences in methods for assessing activity, such as running wheel activity versus spontaneous movement in a home cage, may influence outcomes^22^. Strain differences might also contribute to the variability and make it more difficult to identify these subtle alterations.

Our results support the use of marmosets and mice as non-human models for studying the biological basis of NVP. However, the species differences must be considered carefully^23^. Hormonal dynamics during pregnancy differ across species: marmosets produce mCG instead of hCG^24^, and mice lack chorionic gonadotropin (CG) but secrete placental prolactin-like hormones (PL-I, PL-II)^25^. The timing of estrogen and progesterone rise also varies^23,26^. Moreover, while all three species have hemochorial placentas, the structure of the maternal–fetal barrier differs, with humans having the thinnest, mice the thickest, and marmosets showing an intermediate structure^11,23^. Notably, one proposed cause of NVP, Growth Differentiation Factor 15 (GDF15), is released from the fetus^27,28^, suggesting that the efficiency of molecular transfer from fetus to mother may influence the occurrence or severity of NVP-like phenomena. Actually, the increase in GDF15 levels during pregnancy is lower in mice than in humans and non-human primates^29,31^. While mice remain valuable in many aspects of research, primates may offer advantages due to their greater similarity to humans. Considering that invasive procedures and neural manipulations can be more easily conducted in marmosets than in large primates such as macaques, marmosets may be a particularly practical and promising model for studying the mechanisms of NVP.

Although the present study did not directly investigate the mechanisms underlying NVP, the observation of NVP-like phenomena in both marmosets and mice may provide valuable insight into the possible mechanisms. In both species, the alterations were observed during the stage of placental development, suggesting that NVP-like phenomena may be influenced by placentation-related molecules, such as hCG^2,14,30^. Although mice do not produce CG, other molecules with similar properties may modulate their behavior^25,32^. Further research is needed to elucidate the biological basis of NVP.

In pregnant women, it is difficult to conduct invasive intervention studies due to ethical concerns. The present study provided the possibility for the use of non-human animal models, including marmosets and mice. Using these animal models, we may be able to examine the effectiveness of various methods for conveying NVP. For example, recent studies have revealed that GDF15 is associated with the severity of NVP, making it a potential target for therapeutic intervention^31^. Animal models are essential for testing potential treatments before they can be applied to humans, and our findings may contribute to the development of effective therapies for NVP.

### STAR Methods

#### Marmosets

All experiments using common marmosets were approved by the Animal Experiment Judging Committee of RIKEN (equivalent of Institutional Animal Care and Use Committee, IACUC, approval numbers H28-2-210, H30-2-206, W2020-2-027, H25-2-212, H27-2-212, H29-2-211, W2019-2-011, and W2021-2-033) and were conducted in accordance with the 2011 guideline from the National Research Council of the National Academies. Common marmosets were reared at the RIKEN Center for Brain Science in accordance with the institutional guidelines and under veterinarians’ supervision.

We used 20 adult female marmosets from two different laboratories: Laboratory K (6 mothers, 27 pregnancies) and Laboratory N (14 mothers, 88 pregnancies). Each female was housed in a family cage with her husband and offspring (0-6 individuals per cage). Body weight was measured regularly (Laboratory K: once a week, Laboratory N: once a month) and additionally as needed for health assessments. Water and food were supplied ad libitum. The monkeys’ food was replenished at approximately 11:30, and supplementary foods, such as a piece of sponge cake, dried fruits, and lactobacillus preparation, were given in the afternoon without a fixed schedule. The photoperiod of the colony room was 12L:12D (light period: 8:00-20:00, dark period: 20:00-8:00). Measurements were conducted between 8:00 and 17:00.

#### Clustering analysis

We collected data on the body weight of pregnant females between 150 and 1 day before delivery. Pregnancies were excluded if they had insufficient measurements (i.e., the first measurement was later than 100 days before delivery) or if they resulted in miscarriage or stillbirth. A total of 115 pregnancies (Laboratory K: 27, Laboratory N: 88) were included in the clustering analysis.

Cluster analysis was conducted using R (version 4.4.3)^33^. First, missing body weight values were estimated by linearly connecting the data points to allow analysis of continuous weight patterns. The data from two laboratories were analyzed separately because the frequency of weight measurements differed. To ensure complete data for analysis, we used only the time points for which body weight measurements were available for all pregnancies (Laboratory K: between 121 and 22 days before delivery, Laboratory N: between 112 and 35 days before delivery). The body weight trajectories for each pregnancy were standardized by dividing each data point by the mean weight of that pregnancy. To quantify the pairwise similarity between trajectories, we calculated correlation-based dissimilarities using the *diss* function from the *TSclust* package, with the method set to “COR”^34^. Hierarchical clustering was then performed using the *hclust* function with the default setting. Dendrograms were visually inspected, and clusters were defined by cutting the tree at a height that yielded three major groups. These clusters were named based on cluster size: Cluster A, B, and C in Laboratory K, and as Cluster 1, 2, and 3 in Laboratory N. These clusters were then used to characterize common patterns of weight change during pregnancy.

Reclassification into two superclusters was performed based on the similarity in the two laboratories as follows: Decrease-No and Decrease-Yes. Pregnancies in Decrease-No included Cluster A and Cluster 1, both of which showed continuous body weight increases throughout pregnancies. Decrease-Yes included Cluster B, C, and 2, which showed a transient body weight decrease during mid-pregnancy. Cluster 3, which included only one pregnancy, was classified as “Other”. Data were analyzed between 112 and 35 days before delivery, as all data were available.

#### Mice

All experiments using mice were approved by the Institutional Animal Care and Use Committee of the Faculty of Veterinary Medicine, Hokkaido University (approval numbers 20-0066 and 23-0026). Animals were handled in accordance with the Guide for the Care and Use of Laboratory Animals, Faculty of Veterinary Medicine, Hokkaido University (approved by the Association for Assessment and Accreditation of Laboratory Animal Care International). Water and food were supplied ad libitum. The photoperiod was 12L:12D (light period: 7:00-19:00, dark period: 19:00-7:00).

We used a total of 16 adult C57BL/6JJcl female mice for the experiments. These mice were co-housed with male C57BL/6JJcl mice. The day on which a vaginal plug was observed was designated as gestational day 0 (Gd0), and then the female was singly housed. Pregnant phases were defined as follows. 1^st^ quarter of pregnancy (Q1): from Gd0 to Gd4, 2^nd^ quarter (Q2): from Gd5 to Gd9, 3^rd^ quarter (Q3): from Gd10 to Gd15.

#### Measurement of body weight, food intake, and locomotor activity in mice

Nine pregnant mice were singly housed in standard home cages (170 × 280 × 120 mm). Four food pellets were placed on the perforated top of a box (80 × 120 × 250 mm) and kept in the home cage. At 11:00 each day, the food and box were replaced with new ones, and the body weight of each mouse was measured. Uneaten and spilled food was collected from the box, dried overnight, and weighed. Daily food intake was calculated based on the difference between the provided and recovered food.

Seven pregnant mice were singly housed in special cages (135 × 190 × 160 mm) with an activity sensor (AS-10, Melquest, Japan). Locomotor activity was continuously recorded, and total activity counts were calculated for each 24-hour period.

#### Statistical analysis

Statistical analysis was conducted using R (version 4.4.3)^33^. Cluster analysis in pregnant marmosets is performed as mentioned above. To examine changes in pregnant mice, we used a linear mixed-effects model (LMM) using the *lmer* function from the *lme4* and *lmerTest* packages^35^. Mouse identity was included as a random effect, with random slopes for the gestational day. For food intake, Q2 was included in the model as a binary variable because we hypothesized that Q2 (when placentation progresses) was a physiologically distinct period. In contrast, for locomotor activity, Q1-3 were included in the model as categorical fixed effects (with Q2 as the reference) to evaluate stage-wise changes across pregnancy. Fixed effects included gestational day, pregnancy phase (i.e., Q2 or Q1-3), and their interaction. Model selection was based on Akaike Information Criterion (AIC) using the *dredge* function from the *MuMIn* package^36^.

## Acknowledgement

We thank all staff managing Support Unit for Animal Resources Development in Research Resources Division in the RIKEN Center for Brain Science, the experimental animal facility in the Faculty of Veterinary Medicine at Hokkaido University.

This research was supported by JSPS KAKENHI Grant Number JP19K16901 and JP23K15797 to S.Y.-N., JP18KT0036, JP22K19486, and 22H02664 to K.O.K., RIKEN Center for Brain Science to K.O.K., the Japan Agency of Medical Research and Development (AMED) under grant numbers JP20dm0107144 to K.O.K., Brain/MINDS project 2014 to K.O.K., JST Grant Number JPMJPF2108 to S.Y.-N., Office of Diversity, Equity, and Inclusion at Hokkaido Universit to S.Y.-N., Takeda Science Foundation to K.O.K., Naito Foundation to S. Y.-N., and JSPS J-PEAKS to S.Y.-N.

During the preparation of this work, the authors used ChatGPT and Grammarly in order to improve language and readability. After using these services, the authors reviewed and edited the content as needed and take full responsibility for the content of the publication.

## Author contributions

S.Y.-N. conceived of and organized the study. S.Y.-N., K.S., and T.K. carried out experiments described in Fig. 1-2 under the supervision of K.O.K. Y.S. and K.N. also provided the data for Fig. 1-2. S.Y.-N. and K.K. carried out experiments described in Fig. 3 under the supervision of S.Y. S.Y.-N. analyzed the data with support from S.Q.K., and wrote the manuscript with contributions from all the authors.

